# Stability of Oral and Fecal Microbiome at Room Temperature: Impact on Diversity

**DOI:** 10.1101/2023.11.28.568988

**Authors:** Blanca Rius-Sansalvador, David Bars-Cortina, Olfat Khannous-Lleiffe, Ainhoa Garcia- Serrano, Elisabet Guinó, Ester Saus, Toni Gabaldón, Victor Moreno, Mireia Obón-Santacana

**Affiliations:** Unit of Biomarkers and Susceptibility (UBS), Oncology Data Analytics Program (ODAP), Catalan Institute of Oncology (ICO), L’Hospitalet del Llobregat, 08908 Barcelona, Spain; ONCOBELL Program, Bellvitge Biomedical Research Institute (IDIBELL), L’Hospitalet de Llobregat, 08908 Barcelona, Spain; Doctoral Programme in Biomedicine. University of Barcelona (UB), 08907 Barcelona, Spain; Barcelona Supercomputing Centre (BSC-CNS), 08034 Barcelona, Spain; Institute for Research in Biomedicine (IRB Barcelona), The Barcelona Institute of Science and Technology, 08028 Barcelona, Spain; Consortium for Biomedical Research in Epidemiology and Public Health (CIBERESP), 28029 Madrid, Spain; Catalan Institution for Research and Advanced Studies (ICREA), 08010 Barcelona, Spain; Centro de Investigación Biomédica En Red de Enfermedades Infecciosas (CIBERINFEC), 08028 Barcelona, Spain; Department of Clinical Sciences, Faculty of Medicine and Health Sciences and Universitat de Barcelona Institute of Complex Systems (UBICS), University of Barcelona (UB), L’Hospitalet de Llobregat, 08908 Barcelona, Spain

**Author notes:** Correspondence: Victor Moreno Mireia Obón-Santacana.

**Keywords:** oral microbiome, fecal microbiome, stability study, GCAT, 16S, pilot study

## Abstract

When collecting oral and fecal samples for large epidemiological microbiome studies, optimal storage conditions such as immediate freezing, are not always feasible. It is fundamental to study the impact of temporary room temperature (RT) storage and shipping on the microbiome diversity obtained in different types of samples. We performed a pilot study aimed at validating the sampling protocol based on the viability of the 16S rRNA gene sequencing in microbiome samples.

Fecal and oral samples from five participants were collected and preserved in different conditions: a) 70% ethanol; b) in a FIT tube for stool samples; and c) in a chlorhexidine solution for oral wash samples. Four aliquots were prepared per sample, which were stored at RT, and frozen at days 0, 5, 10 and 15, respectively. In terms of alpha diversity, the maximum average decrease in 5 days was 0.3%, 1.6% and 1.7% for oral, stool in ethanol and stool in FIT, respectively. Furthermore, the relative abundances of the most important phyla and orders remained stable over the two weeks.

The stability of fecal and oral samples for microbiome studies preserved at RT with 70% ethanol, chlorhexidine and in FIT tubes was verified for a 15-day window, with no substantial changes in terms of alpha diversity and relative abundances.

## 1 Introduction

Gut and oral dysbiosis has been associated with the development and progression of some diseases in recent years. For instance, a role of the microbiota has been suggested in an enormous variety of diseases, including metabolic disorders (Bull and Plummer, 2014; Durack and Lynch, 2019; Lu, Xuan and Wang, 2019; Chen, Zhou and Wang, 2021; Fan and Pedersen, 2021; Peng *et al*., 2022), systemic (Willis and Gabaldón, 2020; Martínez *et al*., 2022; Peng *et al*., 2022), cardiovascular (Willis and Gabaldón, 2020; Chen, Zhou and Wang, 2021), liver (Fan and Pedersen, 2021), psychological or mental (Martínez *et al*., 2022), and neurodegenerative diseases (Durack and Lynch, 2019; Chen, Zhou and Wang, 2021; Tuganbaev, Yoshida and Honda, 2022), arthritis (Lu, Xuan and Wang, 2019; Tuganbaev, Yoshida and Honda, 2022), and cancer, such as gastrointestinal cancers (Lu, Xuan and Wang, 2019; Willis and Gabaldón, 2020; Tuganbaev, Yoshida and Honda, 2022), among others.

As many aspects of the relationship between the microbiome and diverse diseases are still unknown (Malla *et al*., 2019), the study of microbiota is an emerging field that is enhancing its knowledge. When collecting samples for microbiome analysis, several procedures and methodologies are used, hindering comparisons across studies. Immediate freezing has been considered the best practice for microbiome preservation (Ilett *et al*., 2019; Moossavi *et al*., 2019; Song *et al*., no date); however, this approach is not feasible for large epidemiological studies that aim to obtain samples shipped by postal mail. In these cases, the samples use to remain for a few days at room temperature until they arrive at their destination (McDonald *et al*., 2018; Williams *et al*., 2019; Young *et al*., 2021; Soriano *et al*., 2022).

Previous research has studied the stability of fecal and oral 16S rRNA gene sequencing microbiome samples (Nechvatal *et al*., 2008; Cardona *et al*., 2012; Carroll *et al*., 2012; Dominianni *et al*., 2014; Choo, Leong and Rogers, 2015; Gorzelak *et al*., 2015; Roberto Flores *et al*., 2015; Tedjo *et al*., 2015; Voigt *et al*., 2015; Sinha *et al*., 2016; Gudra *et al*., 2017; Byrd *et al*., 2019; Bescos *et al*., 2020; Park *et al*., 2020; Krigul *et al*., 2021; Marotz *et al*., 2021; Song *et al*., no date). Regarding fecal microbiome collection methods, 70%-99% ethanol has historically been the most popular stabilization media (Park *et al*., 2020). However, there are fewer studies about the storage of the samples at room temperature compared to other collection methods, such as the Flinders Technology Associates (FTA) or the Fecal Occult Blood Test (FOBT) (Byrd *et al*., 2019).

Furthermore, a widely used collection technique for cancer screening is the Fecal Immunochemical Test (FIT) (Gudra *et al*., 2017). Some metagenomic studies recommend the use of FIT in cohort studies since the microbial profile stability of the samples stored for one week at room temperature has been validated (Gudra *et al*., 2017; Byrd *et al*., 2019; Krigul *et al*., 2021).

The room temperature stability of other fecal microbiome collection methods has been proven for FTA cards at 8 weeks (Song *et al*., no date), OMNIgene Gut Kit for 3 days (Choo, Leong and Rogers, 2015) and 8 weeks (Park *et al*., 2020; Song *et al*., no date), FOBT for 3 (Dominianni *et al*., 2014) and 4 days (Sinha *et al*., 2016; Wu *et al*., 2021) and 1 week (Gudra *et al*., 2017; Byrd *et al*., 2019), RNAlater for 3 (Choo, Leong and Rogers, 2015; Roberto Flores *et al*., 2015), 4 (Sinha *et al*., 2016; Wu *et al*., 2021) and 7 days (Roberto Flores *et al*., 2015; Byrd *et al*., 2019) and 8 weeks (Park *et al*., 2020).

Regarding the oral microbiome, previous studies used Scope® oral wash (mainly composed of Alcohol, Domiphen Bromide and Cetylpyridinium Chloride) to preserve oral microbiome samples (Vogtmann *et al*., 2019; Yano *et al*., 2020), as it has been demonstrated that samples preserved with Scope are stable in terms of alpha and beta diversity up to 4 days at RT (Vogtmann *et al*., 2019; Wu *et al*., 2021). However, as this solution is not easily found in Europe, Chlorhexidine oral wash (Lacer®) was used. Chlorhexidine has been commonly used in many clinical trials where effective results have been proven in reducing the proliferation of bacterial species (Eick *et al*., 2011; James *et al*., 2017; Ben-Knaz Wakshlak, Pedahzur and Avnir, 2019; Brookes *et al*., 2020; Sedghi *et al*., 2021; Xiang, Rojo and Prados-Frutos, 2021). Furthermore, the effect of daily use of chlorhexidine oral wash on the oral microbiome has been studied, showing significant differences in the abundance of some phyla (Bescos *et al*., 2020) and a decrease in terms of alpha diversity compared with sputum samples (Chatzigiannidou *et al*., 2020; Pragman *et al*., 2020). Despite demonstrating that oral washes containing chlorhexidine are related to a major shift in the oral microbiome, the stability of the samples for microbiome analyses, when preserved at RT has not already been studied.

The long-term prospective cohort study of the Genomes for Life (GCAT) aims to facilitate the prediction and treatment of frequent chronic diseases as well as gauge the role of epidemiological, genomic and epigenomic factors (Obón-Santacana *et al*., 2018). In the framework of GCAT, oral and fecal samples for microbiome studies need to be collected throughout the Catalan territory, a northeast region of Spain. Prior to proceeding to its general collection, a validation of the sampling protocol based on the viability of the samples is considered necessary. The main objective of the present study was to investigate the short-term stability at room temperature in both alpha and beta diversity and the distribution of the main bacterial genera of fecal (collected in 70% ethanol and FIT tubes) and oral samples (collected from an oral wash with 0.12% chlorhexidine).

As the samples will be sent by postal mail from different places over the Catalan territory, the logistic challenge regarding the difference in the duration of sample storage at RT is the main point of this study. There is a need to ensure that the quality of the samples in terms of the analysis of microbial diversity is going to be maintained for a few days.

## 2 Materials and Methods

### 2.1 Sample Collection

In this study, 5 volunteer individuals (3 women and 2 men, median age 37) provided three different types of samples for microbiome analysis: one oral wash, preserved in 0.12% chlorhexidine and two fecal samples, one preserved in a FIT tube (FIT, OCSensor, Eiken Chemical Co., Tokyo, Japan) and another in a 5 ml tube with 1 ml of 70% ethanol. Samples were collected at home. Participants were instructed to obtain oral samples in the morning, before any food or tooth brushing, by doing an oral rinse for 1 minute with Lacer® oral wash and then spitting the content in a tube. Stool samples, if obtained the night before, were kept at 4°C before transport to the lab. For the three collection methods, a total of 4 aliquots of each sample were prepared and one aliquot was immediately frozen at -80°C until processing. The rest were consecutively frozen after remaining at room temperature for 5, 10 and 15 days, resulting in a total of 60 samples from 5 individuals at 4 time points (**Figure 1; Supplementary Table S1**). None of the participants took oral antibiotics, injected antibiotics, stomach protectors, or acid-lowering medication in the last 3 months. All individuals agreed to participate in the study and provided written informed consent. The University Hospital of Bellvitge ethics committee approved the protocol of the study (PR084/16).

**Figure 1.**
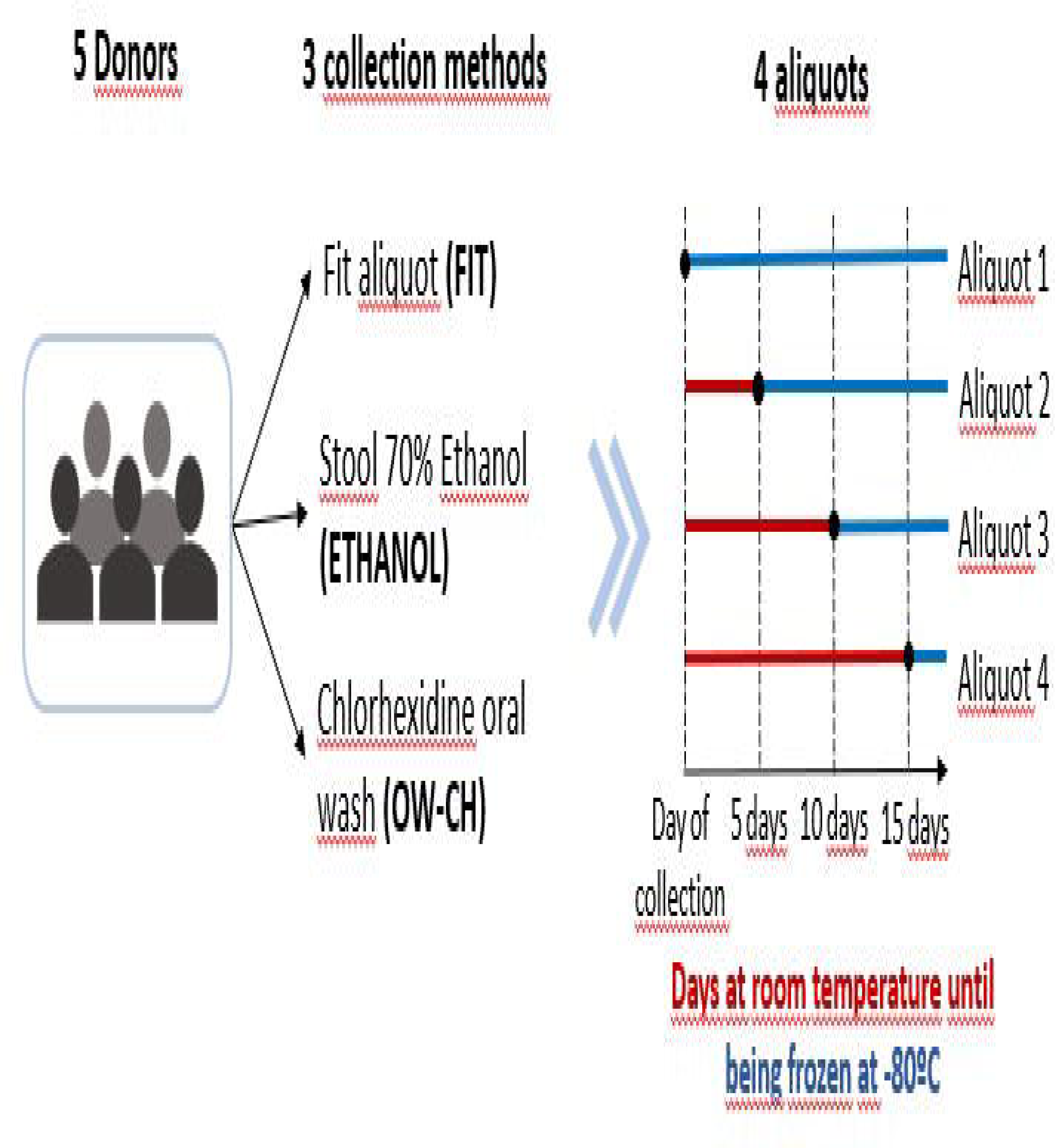
Sample collection diagram

### 2.2 DNA Extraction and Sequencing

DNA was extracted using the DNeasy PowerLyzer PowerSoil Kit (Qiagen, ref. QIA12855) following the manufacturer’s instructions with slight modifications depending on the initial sample type (FIT, oral wash and stool samples). Two negative controls of the DNA extraction process (with no initial sample) were also included. Briefly, for FIT samples, a pre-enrichment step was added by centrifuging the samples at 20,000 g for 5 min at 4°C. The supernatant was removed, and the pellet was resuspended in 750 μl of PowerBead Solution, mixed and transferred to a Bead tube with beads. Stool samples were already frozen in 2 ml tubes, where 750 μl of PowerBead Solution and the beads were directly added. For oral wash samples, pellets were resuspended in 750 μl of PowerBead Solution, mixed and transferred to a Bead tube with beads. From here on all samples were processed in the same way: 60 μl of Solution C1 was added, and samples were vortexed briefly and incubated at 70°C with shaking (700 rpm) for 10 min. The extraction tubes were then agitated in a horizontal vortex (Genie2) for 20 min at maximum speed. Tubes were centrifuged at 10,000 g for 3 min and the supernatant was transferred to a clean tube. Then, 250 μl of Solution C2 was added, and the samples were vortexed for 5 s and incubated on ice for 5 min. After 1 min of centrifugation at 10,000 g, 600 μl of the supernatant was transferred to a clean tube, 200 μl of Solution C3 was added, and the samples were vortexed for 5 s and incubated on ice for 5 min again. A total of 750 μl of the supernatant was transferred into a clean tube after 1 min centrifugation at 10,000 g. Then, 1,200 μl of Solution C4 was added to the supernatant, samples were blended by pipetting up and down, and 675 μl was loaded onto a spin column and centrifuged at 10,000 g for 1 min, discarding the flow through. This step was repeated three times until all samples had passed through the column. 500 μl of Solution C5 was added onto the column, and the samples were centrifuged at 10,000 g for 1 min. The flow through was discarded, and one extra minute of centrifugation at 10,000 g was performed to dry the column. Finally, the column was placed into a new 2 ml tube for final elution with 50 μl of Solution C6 and centrifugation at 10,000 g during 30s. For DNA quality control, two serial dilutions of the DNA samples were used. Genomic DNA was quantified using SYBRGreen I (Sigma‒Aldrich, Merck) and the total bacterial load in the DNA sample was estimated by a real-time PCR assay with primers described in Nadkarni et al. 2002 (Nadkarni *et al*., 2002) (forward 5’-TCCTACGGGAGGCAGCAGT-3’ and reverse primer 5’-GGACTACCAGGGTATCTAATCCTGTT-3’), using a 7900 HT Fast Real-Time PCR System (Applied Biosystems).

For library preparation, the DNA samples were normalized according to their bacterial DNA content to be used as a template to prepare 16S rRNA libraries (region V3-V4). The 16S rRNA V3-V4 region was amplified with primers previously described (Willis *et al*., 2018), but the library preparation protocol was slightly modified. First, normalized DNA samples were used to amplify the V3–V4 regions of the 16S ribosomal RNA gene, in a limited cycle PCR. The PCR was performed in a 25 μl volume with 0.08 μM primer concentration and NEBNext Q5 Hot Start HiFi PCR Master Mix (ref. M0543L, New England Biolabs). The cycling conditions were an initial denaturation of 30 s at 98°C followed by 5 cycles of 98°C for 10 s, 55°C for 5 min, and 65°C for 45 s. After this first PCR, a second PCR was performed in a total volume of 50 μl. The reactions comprised NEBNext Q5 Hot Start HiFi PCR Master Mix and Nextera XT v2 adaptor primers. PCR was carried out to add full-length Nextera adapters: initial denaturation of 30 s at 98°C followed by 17 cycles of 98°C for 10 s, 55°C for 30 s, and 65°C for 45 s, ending with a final elongation step of 5 min at 65°C. Libraries were purified using AgenCourt AMPure XP beads (ref. A63882, Beckman Coulter) with a 0.9X ratio according to the manufacturer’s instructions and were analyzed using Fragment Analyzer (ref. DNF-915, Agilent Biosystems) to estimate the quantity and check size distribution. A pool of normalized libraries was prepared for subsequent sequencing. Final pools were quantified by qPCR using the Kapa library quantification kit for Illumina Platforms (Kapa Biosystems) on an ABI 7900HT real-time cycler (Applied Biosystems). Sequencing was performed on an Illumina MiSeq with 2 × 300 bp reads using v3 chemistry with a loading concentration of 18 pM. To increase the diversity of the sequenced, 10% of PhIX control libraries were spiked in.

Negative controls of the PCR amplification steps were routinely performed in parallel using the same conditions and reagents. Our control samples systematically provided no visible band or quantifiable DNA amounts. The ZymoBIOMICS™ Microbial Community DNA Standard (ref. D6306, Zymo) was amplified and sequenced in the same manner as all other experimental samples.

### 2.3 Bioinformatics and statistical analysis

Raw data were processed by using the Dada2 pipeline (v. 1.12.1) (Callahan *et al*., 2015). Low-quality reads were filtered and trimmed out based on the observed quality profiles by using the *filterAndTrim* function, truncating forward and reverse reads below 290 and 230, respectively, and considering a value of 2 as the maximum expected error. Furthermore, 10 reads from the start of each read were removed. We combined identical sequencing reads into unique sequences, made a sample inference from a matrix of estimated learning errors and merged paired reads. Chimeric sequences were removed by using the *removeBimeraDenovo* function, and taxonomy was assigned utilizing the SILVA 16S rRNA database (v.138) (Quast *et al*., 2013).

Two negative controls from DNA extraction were analyzed to assess possible sources of contamination and removed for further analysis. The resulting Amplicon Sequence Variant (ASV) table was merged with the metadata creating a *phyloseq* (v. 1.26.1) (McMurdie and Holmes, 2013) object. We filtered out taxa with fewer than 100 reads and with a relative abundance less than 0.1% or present in less than 5% of the samples.

Statistical analyses were performed using the 4.1.2 R version. In order to adjust for differences in the number of reads across samples and allow a proper alpha diversity comparison (Willis, 2019), the data were sampled at a value of 42,321 (rarefaction efficiency index = 0.99 (Hong *et al*., 2022), the minimum number of reads (**Supplementary Figure S2**).

To assess the alpha diversity of the samples four indexes were calculated (Thukral, 2017; Datta and Guha, 2021) (Chao, Simpson, Inverse Simpson and Shannon). However, since analogous results were obtained, only the Shannon index is reported in this study, which considers the differences in the abundance of each species and is the most commonly used diversity metric (Reese and Dunn, 2018). In addition, the mean and range richness of the samples at all taxonomical levels were plotted for each time point. Furthermore, the OTUs found in immediately frozen samples and not found anymore were listed.

For the purpose of studying beta diversity, Bray‒Curtis, Jaccard, unweighted UniFrac and weighted UniFrac dissimilarity distances were considered (Plantinga and Wu, 2021; By IMPACTT investigators, 2022), but since similar results were obtained, only the Bray‒Curtis dissimilarity is reported. The projections of the individuals were plotted in one of the three plots depending on the collection method (**Figure 2**). Furthermore, the shapes were plotted according to the days staying at room temperature and colored according to the sample number.

**Figure 2.**
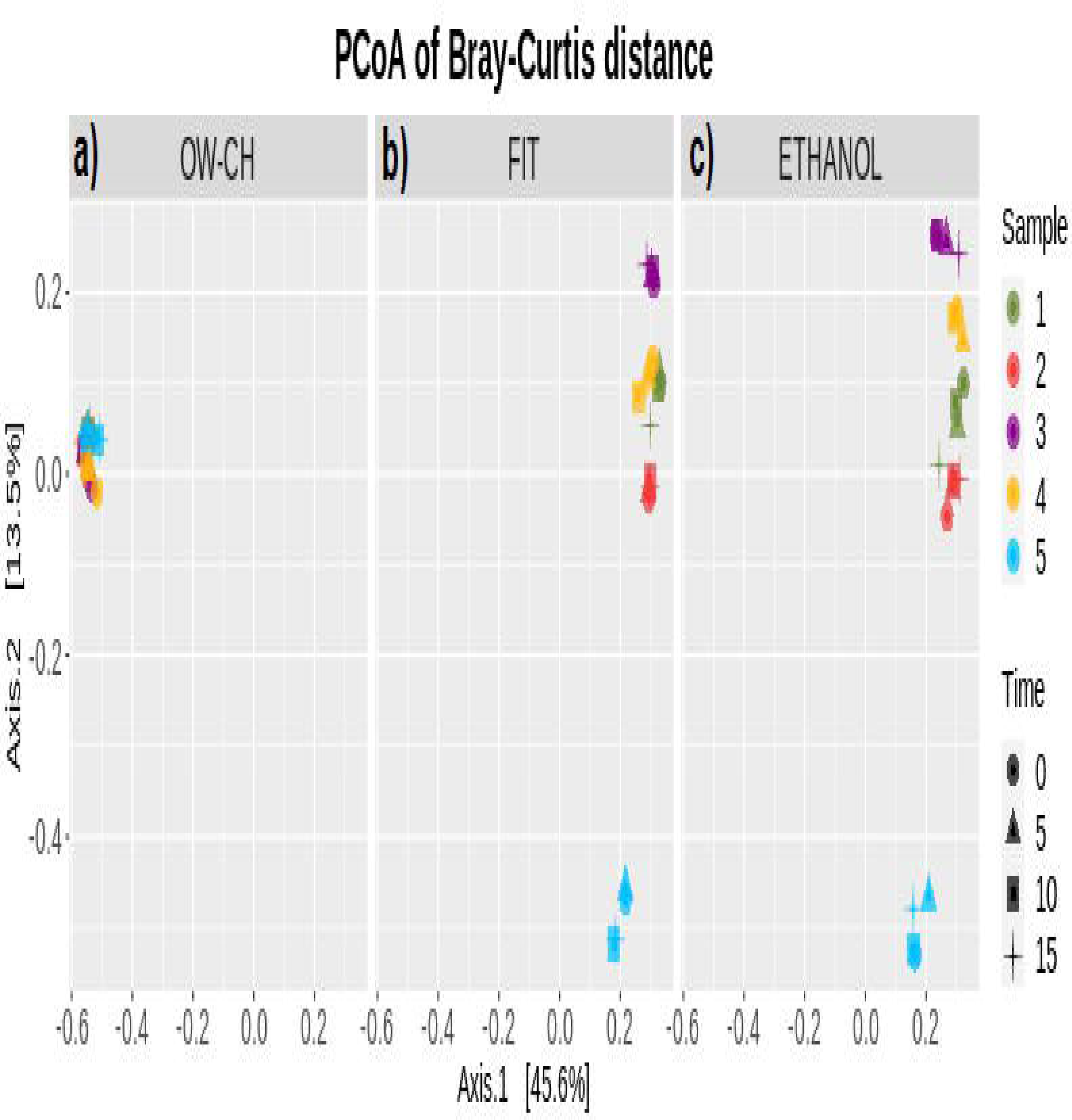
Principal Coordinates Analysis based on Bray‒Curtis dissimilarity matrix stratified by method, sample and days at room temperature representing beta diversity

Since the sample size of this stability study was small, the statistical analysis was focused on the estimation of changes and their 95% confidence intervals. Linear mixed models (LMM) were used to estimate the change in alpha diversity over the time points 0, 5, 10 and 15 days. LMMs account for the correlations between data including the subject as a random effect. Estimated marginal means (EMMs) were used to estimate differences among time points.

Multiple analysis of variance (MANOVA) was conducted to compare the abundance of the top 5 phyla and the top 20 orders with the number of days that the sample remained at room temperature before being frozen.

A sensitivity analysis was performed, removing one subject that showed a pattern considerably different from others.

The dataset that was generated and analyzed in our study is available at the Zenodo repository (DOI: 10.5281/zenodo.7684999, accessed on 28th February 2023).

## 3 Results

### 3.1 Comparing Alpha and Beta Diversity between Methods

The 70% ethanol and FIT collection methods for stool showed small differences in terms of the Shannon index of diversity at time 0 (difference = 0.23, 95% CI 0.18 to 0.65). We observed a larger dispersion of diversity values for 70% ethanol. The alpha diversity of stool samples measured by the Shannon index was similar to that of oral wash (OW-CH), although these samples had a different overall composition (**Figure 3**).

**Figure 3.**
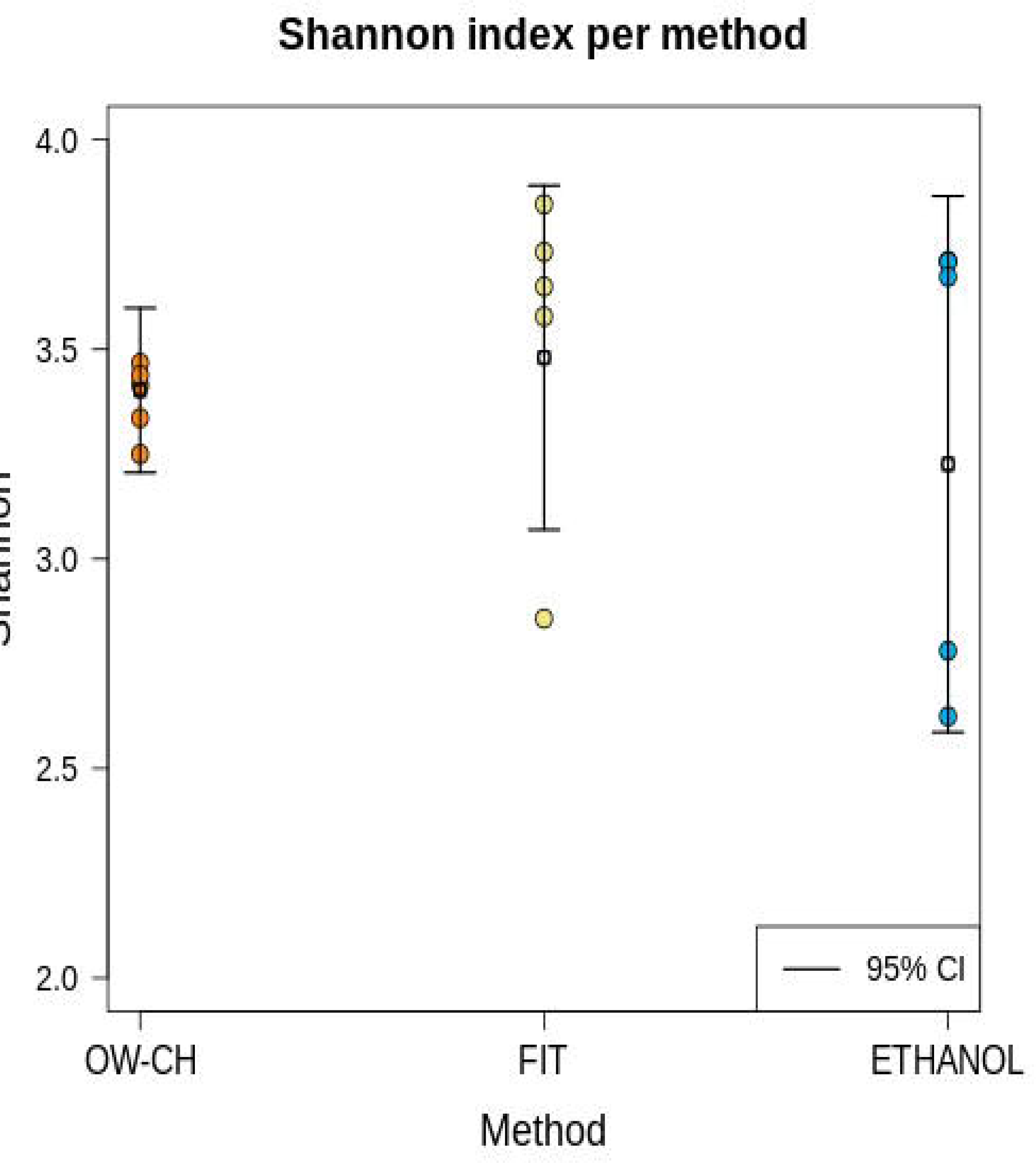
Shannon index plot for each method in immediately frozen samples, its mean and 95% Confidence Interval

In terms of the richness of the OTUs it was not possible to see a significant decrease or trend among the days at room temperature (**Figure 4**). Nevertheless, we found a few OTUs present in the immediately frozen samples that were no longer present in the samples stored at room temperature (**Supplementary Table S3)**.

**Figure 4.**
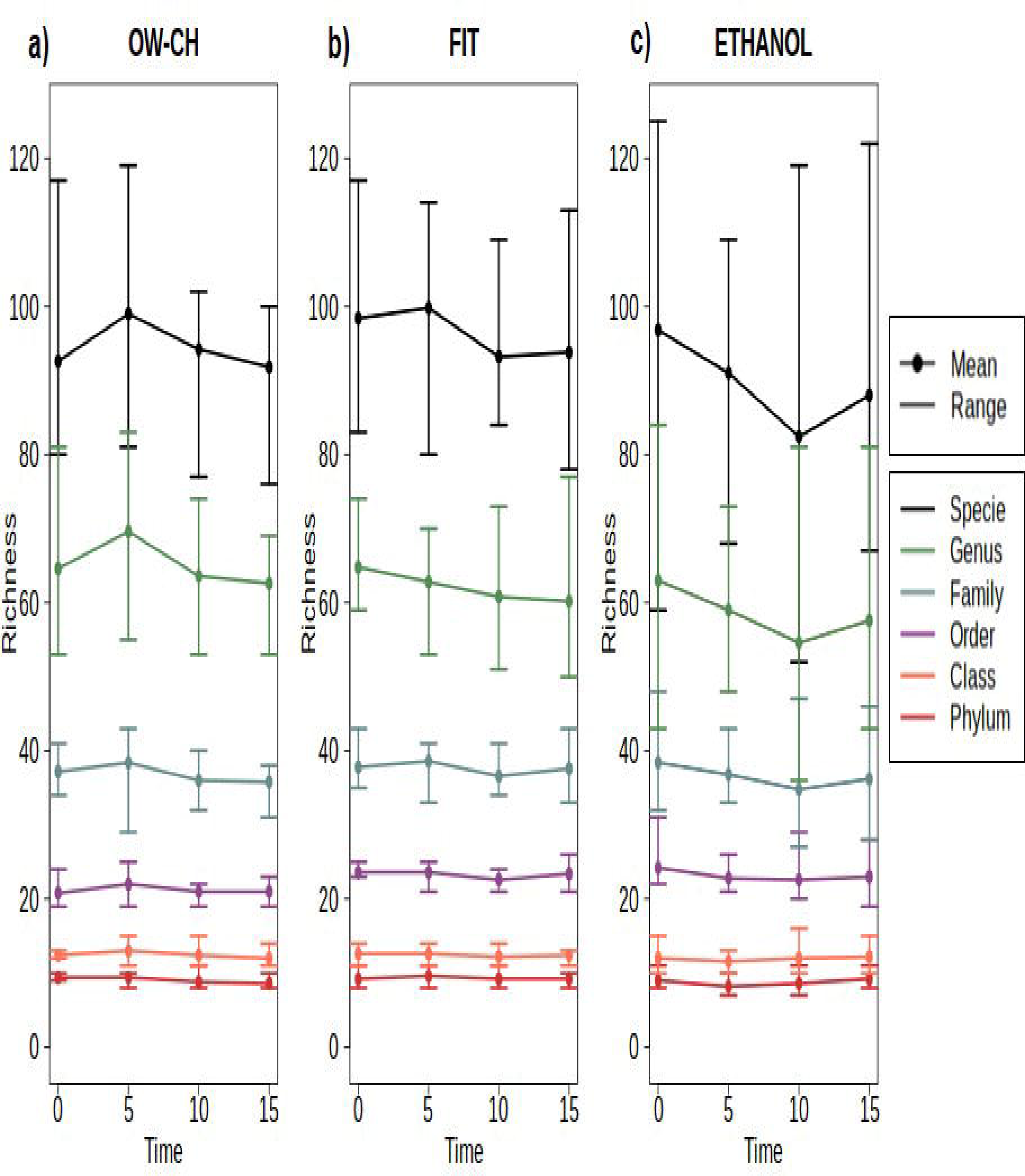
Mean and range richness of the samples at all taxonomy levels among the days at room temperature for each sequencing method

Regarding beta diversity, it can be noticed that oral microbiota (**Figure** ¡Error! No se encuentra el origen de la referencia.**2a)** has a different distribution from stool microbiota (**Figure 2b** and ¡Error! No se encuentra el origen de la referencia.**Figure 2c**), but both preservation methods for stool samples show very good agreement when accounting for Bray‒Curtis dissimilarity distance. In addition, it can be observed that the projections of the individuals are grouped by individual, not by time. Therefore, there cannot be observed differences or patterns to do with the days stored at room temperature. On the second axis, subject 5 was more distant than others for stool samples. This subject also showed lower alpha diversity (**Figure 5**).

**Figure 5.**
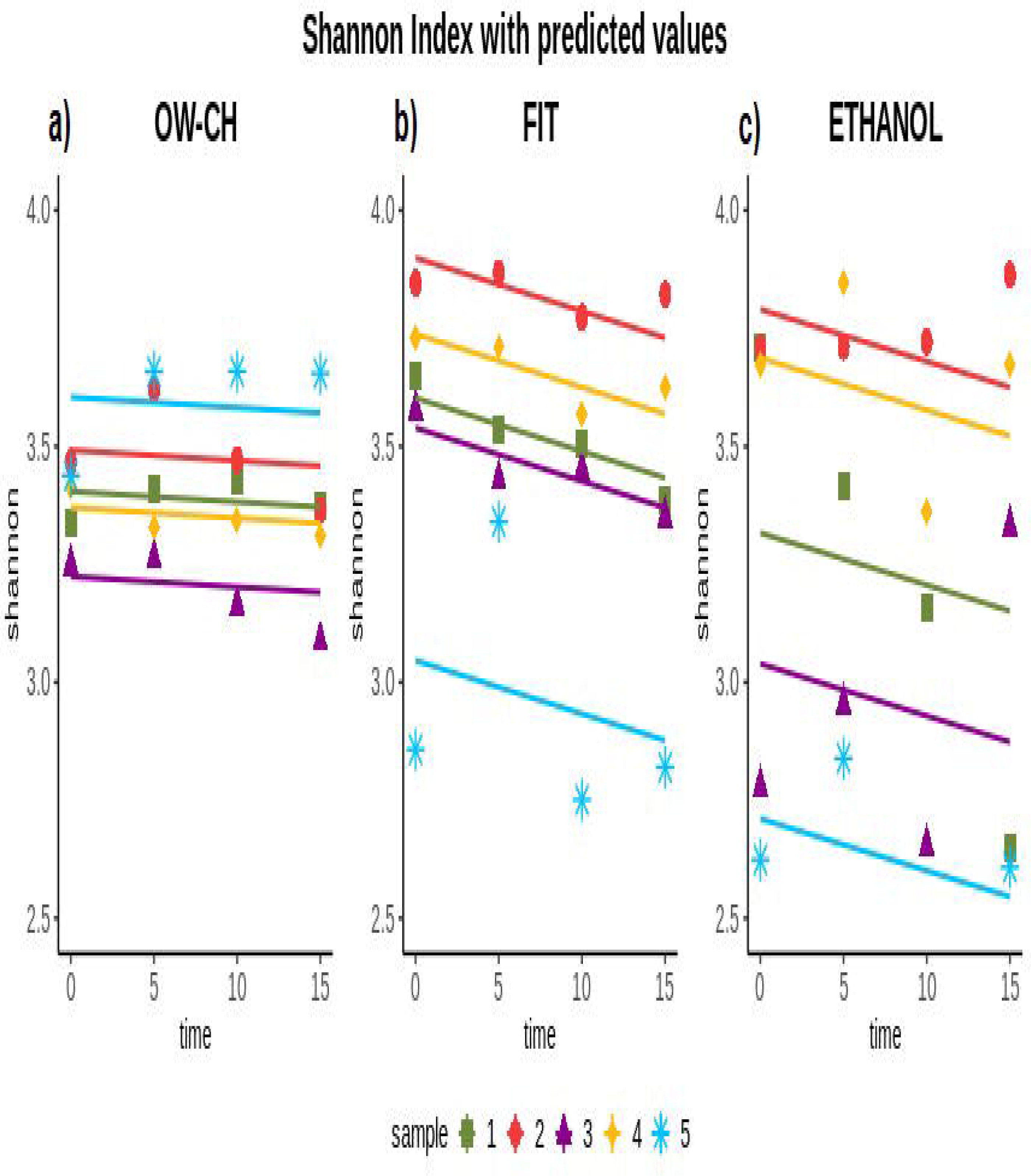
Shannon index at each time point and predicted values based on Generalized Linear Mixed Models for the three methods

**Figure 6.**
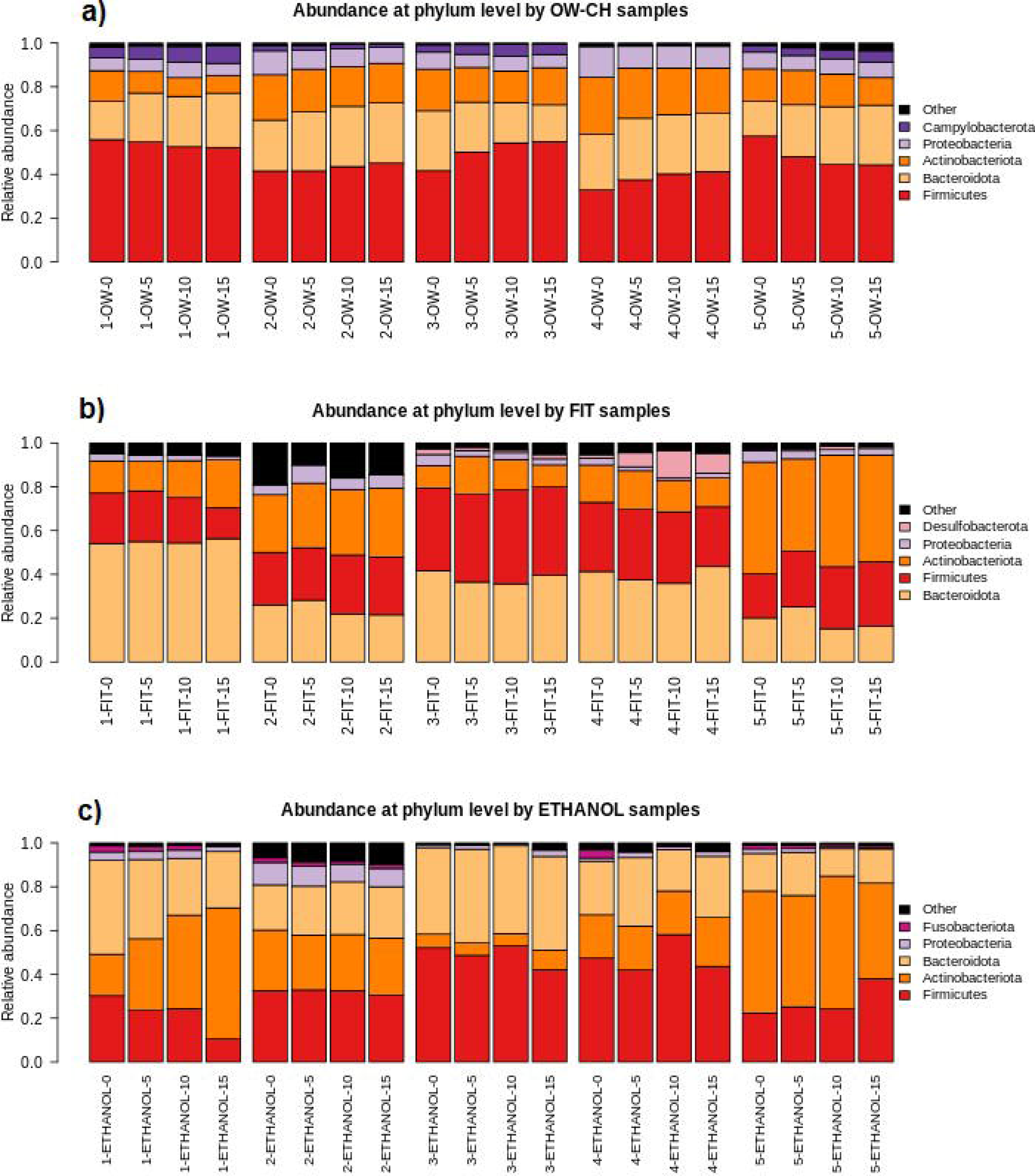
Relative abundance plots for the main phyla per method and individual among the time at room temperature

## 3.2 Stability of the Samples

The Shannon index for oral samples preserved in chlorhexidine was stable over the 15 days at RT, with no substantial trend (**Table 1**; **¡Error!** No se encuentra el origen de la referencia. 5). However, the alpha diversity index decreased over time for stool samples. Nevertheless, although the shipment of the samples is not expected to take so long, the Shannon Index variation over the 15 days was only - 3.68% for the FIT collection method (¡Error! No se encuentra el origen de la referencia.). For the 70% ethanol and oral wash collection methods, the shifts were -2.12% and -0.59%, respectively. The pairwise comparison of time points 0 and 5 did not show a major change.

**Table 1.** Slopes of the Generalized Linear Mixed Models for Shannon Index with 95% confidence intervals and 5-day percentage mean decrease.

| Shannon Index |  |  |  |  |
| --- | --- | --- | --- | --- |
| Method | Average time 0 | Slope coefficient | 95% CI | 5-day % decrease |
| OW-CH | 3.38 | -0.002 | (-0.009; 0.005) | -0.32 |
| FIT | 3.53 | -0.011 | (-0.021; -0.001) | -1.58 |
| ETHANOL | 3.30 | -0.011 | (-0.034; 0.012) | -1.67 |

**Table 2.**
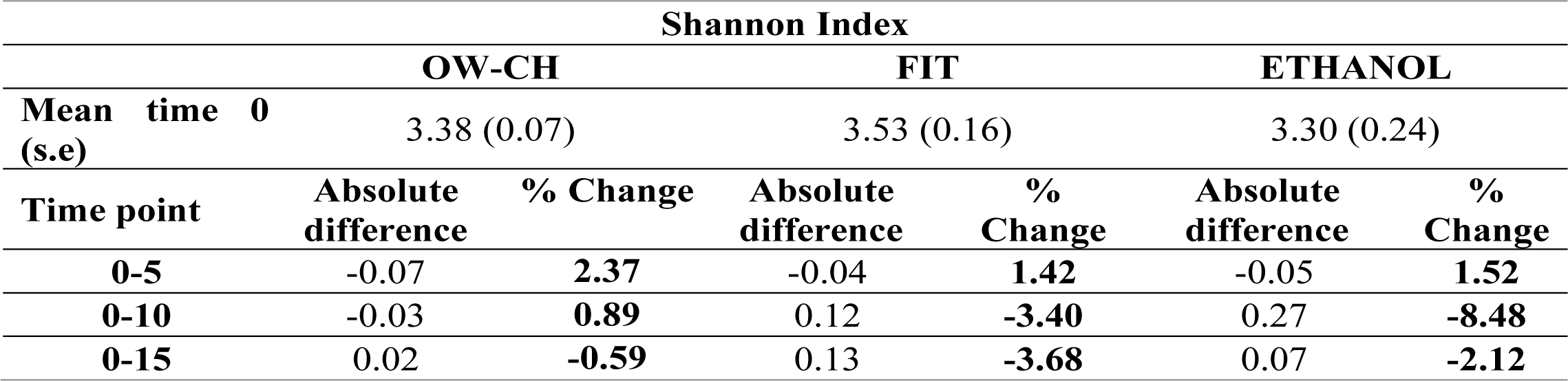
Mean at time 0 (standard error) and absolute difference and percentage of change of pairwise comparisons of the time points with respect to day 0. Average values were derived from the estimated marginal means of the LMM model.

The results remained unchanged when we performed the sensitivity analysis excluding subject 5 since it was found apart from the rest in the Principal Coordinates Analysis plot, both with Jaccard and Bray‒ Curtis dissimilarity matrices.

## 3.3 Top 5 Phyla Stability over Time

The results from the MANOVA (**Supplementary Table S4**) indicated no differences regarding the days that the samples were stored at room temperature in the relative abundances of the most common phyla (*Firmicutes*, *Bacteroidota*, *Actinobacteriota*, *Proteobacteria* and *Campylobacterota*)

## 3.4 Top 20 Orders Stability over Time

MANOVA comparisons between the top 20 orders and the days at room temperature (**Supplementary Table S5**) only showed differences in a few low-frequency orders. At 15 days, *Burkholderiales* decreased by 7.8% and 15.1% in the oral wash and FIT samples, respectively, and *Synergustales* decreased by 57.3% in the FIT samples. No major shifts were found.

The fecal microbiome at the order level of sample 5 was different from the others; however, it showed similar stability patterns (**Figure 7**).

**Figure 7.**
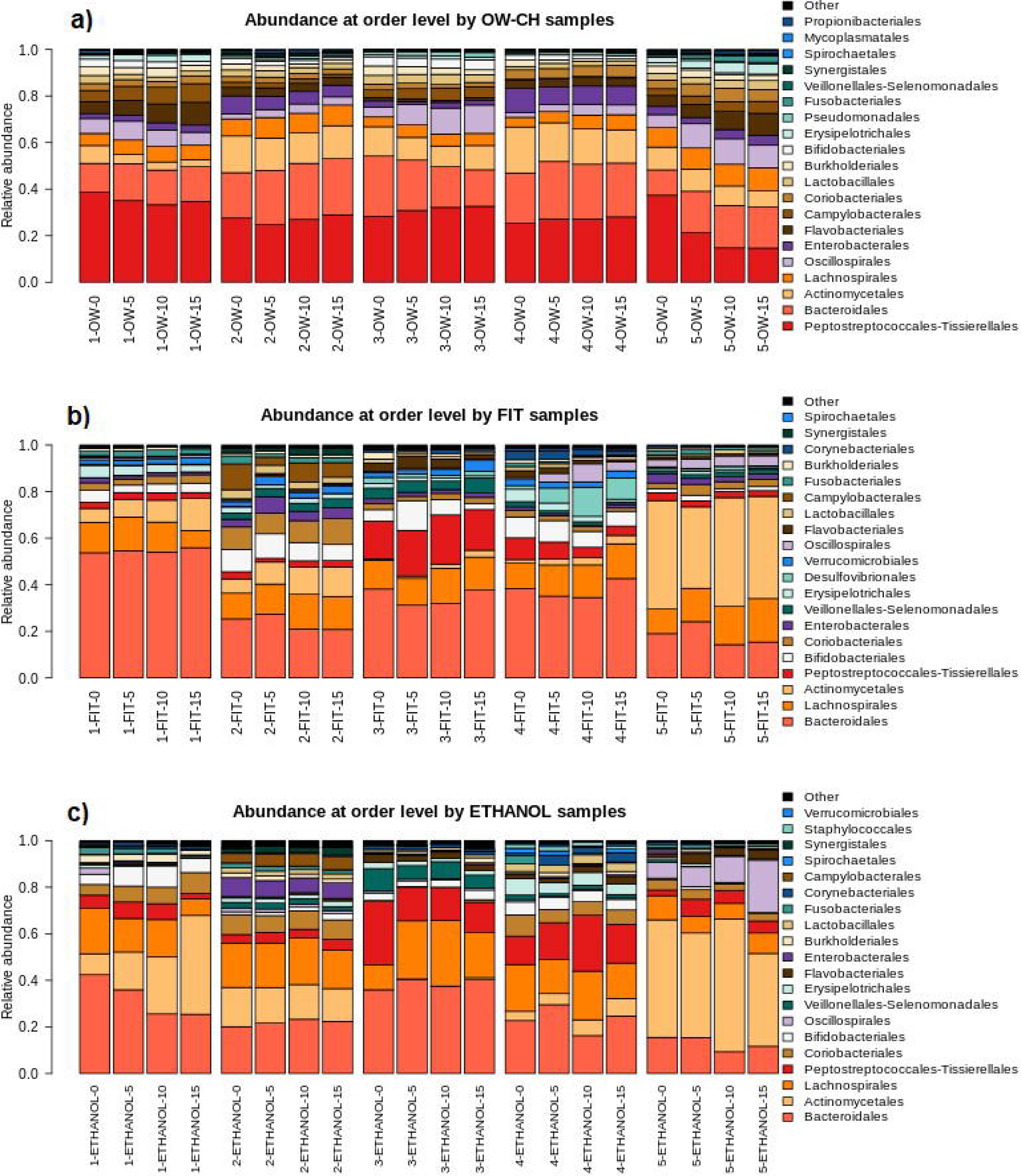
Relative abundance plots for the main order per method and individual among the time at room temperature

## 4 Discussion

The stability of microbiome samples at room temperature for 15 days was investigated for oral wash samples preserved in chlorohexidine and two fecal collection methods (FIT and 70% ethanol). We found that oral microbiome diversity and composition were, in general, very stable during the 15 days at room temperature. For both fecal preservation methods, however, a small decrease in diversity was observed, mainly after day 5, with the samples stored in ethanol showing more heterogeneity. Between subjects, variability was of similar magnitude to the fluctuations in alpha diversity observed over time.

Although the microbial profile stability has previously been validated for 95% ethanol (Byrd *et al*., 2019; Marotz *et al*., 2021; Song *et al*., no date), several studies caution against the use of 70% ethanol since it is found to be less stable than other collection methods stored for 4 days (Sinha *et al*., 2016; Byrd *et al*., 2019), 1 week (Sinha *et al*., 2016; Marotz *et al*., 2021) and 8 weeks (Song *et al*., no date). Other works do not report significant changes between immediately frozen samples and the 70% ethanol microbiome samples stored for 8 weeks at room temperature (Park *et al*., 2020). Our results are in agreement with previous works that reported no significant changes between immediately frozen samples and 70% ethanol samples for microbiome studies, at least for 5 days at room temperature, which is the usual shipment time.

Regarding the FIT collection method, several studies recommend its use in epidemiological studies. It has been proved that, in terms of alpha diversity, FIT samples remain stable for one week at RT (Gudra *et al*., 2017; Byrd *et al*., 2019; Krigul *et al*., 2021). Our work agrees with previous research, not showing significant changes in the composition of the samples.

Previous studies used Scope® oral wash to preserve oral microbiome samples (Vogtmann *et al*., 2019; Yano *et al*., 2020), and its stability at room temperature was already verified (Vogtmann *et al*., 2019; Wu *et al*., 2021); however, it is not easily found in Europe. As chlorhexidine has been commonly used in many clinical trials where effective results have been proven in reducing the proliferation of bacterial species (Eick *et al*., 2011; James *et al*., 2017; Ben-Knaz Wakshlak, Pedahzur and Avnir, 2019; Brookes *et al*., 2020; Sedghi *et al*., 2021; Xiang, Rojo and Prados-Frutos, 2021), we opted for Lacer® Chlorhexidine oral wash to preserve the samples. To the best of our knowledge, the stability of Lacer® oral wash samples at room temperature has not been previously studied. Our results sustain that the alpha diversity of the samples remained stable for 15 days at RT with no major shifts.

Although we are aware that the small sample size of the present study is not powered to perform statistical tests, the estimates of change and 95% confidence intervals allow a reasonable assessment of the quality of the sample preservation methods. On average, the magnitude of the changes in alpha diversity was smaller than 2%, allowing a reasonable assessment of the quality of the sample preservation methods. Phylum compositions showed good temporal stability, except for fecal samples preserved in ethanol in subject number 5, which had a different microbiome pattern. Furthermore, a shift was observed in individual 1 for the 70% ethanol samples, while *Actinobacteriota* increased and *Firmicutes* and *Bacteroidota* decreased. Regarding order compositions, although slight relative abundance differences could be found in a few of the low-abundance orders, the main ones remained stable during the time of the study.

## 5 Conclusion

To conclude, the stability of the samples regarding diversity and composition was verified for the chlorohexidine oral wash and two fecal methods (FIT and 70% ethanol). Alpha diversity was maintained over 15 days at room temperature for the chlorohexidine oral wash. For fecal samples, both 70% ethanol and FIT showed a decrease in diversity over time but a small decrease during the first 5 days. The relative abundance of the top 5 phyla and the top 20 orders was verified to be consistent for the three methods.

## 6 Author Contributions

VM, MOS and EG conceptualization. VM, MOS, EG and BRS data curation. VM, MOS, BRS, AGS, OKL, ES and TG formal analysis. BRS, MOS and VM writing – original draft. DBC, AGS, EG, OKL, ES and TG Writing – review & editing. All authors have read and agreed to the published version of the manuscript.

## 8 Acknowledgments

We gratefully thank all volunteers and staff for their time and commitment to the study. The Lacer company freely provided Chlorhexidine mouthwash to carry out this study. The authors thank the CERCA Program/Generalitat de Catalunya for their institutional support and the Genomics Unit at the CRG (Centre for Genomic Regulation, Barcelona, Spain) for assistance with 16S sequencing.

## 10 Data Availability Statement

The dataset supporting the conclusions of this article is available in the Zenodo repository, https://zenodo.org/records/7684999 (DOI: c, accessed on 28th February 2023). Raw data is available in ENA with project accession PRJEB67775 (https://www.ebi.ac.uk/ena/browser/view/PRJEB67775), accessed on 27^th^ October 2023.

## 12 Ethics declarations

No animal studies are presented in this manuscript.

The studies involving humans were approved by University Hospital of Bellvitge (PR084/16). The studies were conducted in accordance with the local legislation and institutional requirements. The participants provided their written informed consent to participate in this study.

No potentially identifiable images or data are presented in this study.

