## Supplemental Table S1 for "Stability of Oral and Fecal Microbiome at Room Temperature: Impact on Diversity"

**Additional Table S1** Sample size and collection methods for oral and faecal alpha diversity microbiome analysis

|  | n samples frozen at different timepoints | | | | |
| --- | --- | --- | --- | --- | --- |
| Type of samples | Day 0 | Day 5 | Day 10 | Day 15 | Total |
| Oral Sample Collection Method |  |  |  |  |  |
| OW-CH | 5 | 5 | 5 | 5 | 20 |
| Faecal Sample Collection Method |  |  |  |  |  |
| FIT | 5 | 5 | 5 | 5 | 20 |
| ETHANOL | 5 | 5 | 5 | 5 | 20 |
| Total | 15 | 15 | 15 | 15 | **60** |
