## Supplemental Figure S2 for "Stability of Oral and Fecal Microbiome at Room Temperature: Impact on Diversity"

**Additional Figure S2** Rarefaction curves for each alpha diversity index; in red the minimum depth

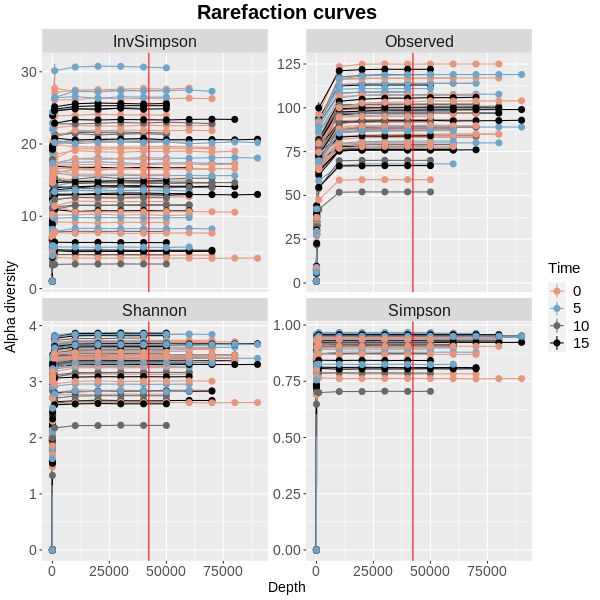
