## Supplemental Table S3 for "Stability of Oral and Fecal Microbiome at Room Temperature: Impact on Diversity"

**Additional Table S3** OTUs present in the immediately frozen samples and not anymore in the room temperature stored ones.

|  | | OW-CH | FIT | ETHANOL |
| --- | --- | --- | --- | --- |
| Phylum | | - | - | Patescibacteria  Unclassified P2 |
| Class | | **-** | Lentisphaeria  Vampirivibrionia | Gracilibacteria  Saccharimonadia  Unclassified C3 |
| Order | | Monoglobales  Unclassified O14 | Acidaminococcales  Clostridia UCG 014  Coxiellales  Gastranaerophilales  RF39 | Clostridia vadinBB60 group  JGI 0000069 P22  Opitutales  Peptococcales  RF39  Saccharimonadales  Unclassified O11 |
| Family | | Aerococcaceae  Burkholderiaceae  Defluviitaleaceae  Monoglobaceae  Unclassified F109  Unclassified F29 | Acidaminococcaceae  Coxiellaceae  Hydrogenoanaerobacterium  Unclassified F112  Unclassified F114  Unclassified F115  Victivallaceae | Peptococcaceae  Puniceicoccaceae  Saccharimonadaceae  UCG 010  Unclassified F114  Unclassified F116  Unclassified F119  Unclassified F22 |
| Genus | | Abiotrophia  Agathobacter  Amnipila  Anaerofilum  Butyricicoccus  Catenibacterium  Centipeda  Defluviitaleaceae UCG 011  Family XIII AD3011 group  Fusicatenibacter  Lachnospiraceae UCG 003  Lachnospiraceae UCG 004  Lautropia  Monoglobus  Sutterella  Unclassified G255  Unclassified G267  Unclassified G62  Veillonella | Eubacterium ruminantium group  Aggregatibacter  CAG 56  Christensenellaceae R 7 group  Colidextribacter  Coxiella  Eggerthia  Eikenella  Flavonifractor  Incertae Sedis  Intestinimonas  Lachnospiraceae AC2044 group  Lachnospiraceae ND3007 group  Lachnospiraceae NK4A136 group  Limosilactobacillus  Marvinbryantia  Phascolarctobacterium  Streptobacillus  Subdoligranulum  TM7x  UCG 004  Unclassified G258  Unclassified G261  Unclassified G263  Unclassified G268  Unclassified G269  Unclassified G281  Victivallis | Eubacterium brachy group  Allisonella  CAG 56  Candidatus.Saccharimonas  Eggerthia  Flavonifractor  Intestinibacter  Lachnospiraceae AC2044 group  Lachnospiraceae ND3007 group  Lachnospiraceae UCG 004  Levilactobacillus  Limosilactobacillus  Mailhella  Marvinbryantia  Parvimonas  Prevotellaceae UCG 001  Rikenellaceae RC9 gut group  Streptobacillus  Unclassified G261  Unclassified G264  Unclassified G271  Unclassified G273  Unclassified G278  Unclassified G282  Unclassified G32  Unclassified G45 |
| Specie | Neisseria subflava  Olsenella unclassified S1181  Prevotella histicola  Prevotella nigrescens  Capnocytophaga leadbetteri  Simonsiella unclassified S1366  Streptococcus constellatus  Streptococcus anginosus  Capnocytophaga unclassified S1216  Treponema refringens  Fusobacterium unclassified S1352  Unclassified unclassified S1348  Unclassified unclassified S160  Leptotrichia buccalis  Simonsiella muelleri  Prevotella oulorum  Treponema unclassified S1374  Prevotella veroralis  Shuttleworthia satelles  Eikenella unclassified S1363  Eggerthia catenaformis  Fretibacterium unclassified S1376  Kingella unclassified S1364  Prevotella loescheii  Comamonas unclassified S1361  Scardovia wiggsiae  Aestuariimicrobium unclassified S1178 | | Streptococcus oralis  Veillonella atypica  Streptococcus unclassified S55  Streptococcus australis  Roseburia hominis  Unclassified unclassified S1379  Unclassified unclassified S1218  Escherichia Shigella unclassified S1370  Roseburia intestinalis  Lachnospiraceae UCG 001 bacterium  Enterobacter hormaechei  Eubacterium hallii group unclassified S1257  Lactobacillus helveticus  Ruminococcus flavefaciens  Bacteroides thetaiotaomicron  Unclassified unclassified S1239  Lachnospiraceae UCG 003 unclassified S1287  Erysipelotrichaceae UCG 006 unclassified S1233  Lachnospiraceae FCS020 group unclassified S1282  DNF00809 unclassified S1189  Negativibacillus unclassified S1321  Megamonas unclassified S1346  Odoribacter unclassified S1201  Unclassified unclassified S1328  Oscillospira unclassified S1308  Lactobacillus iners  Lachnospiraceae AC2044 groupunclassified S1281  Fenollaria unclassified S1338  Coprobacter secundus  GCA 900066575 unclassified S1275  Enorma unclassified S1184  Allisonella unclassified S1349  Paraprevotella clara  Catenibacillus unclassified S1269  Turicibacter sanguinis  UCG 008 unclassified S1299  UCG 009 unclassified S1300 | Streptococcus oralis  TM7x unclassified S1359  Veillonella dispar  Rothia mucilaginosa  Streptococcus australis  Unclassified unclassified S351  Lactobacillus unclassified S202  Eggerthella unclassified S1190  Bifidobacterium bifidum  Intestinimonas unclassified S1305  Enterorhabdus unclassified S1191  Unclassified unclassified S1302  Ruminococcus flavefaciens  Bacteroides thetaiotaomicron  Propionibacterium freudenreichii  Streptococcus mutans  Lachnospiraceae FCS020 group unclassified S1282  Olsenella scatoligenes  DNF00809 unclassified S1189 Negativibacillus unclassified S1321  unclassified unclassified S160  Megamonas unclassified S1346 Oscillospira unclassified S1308  Unclassified unclassified S1199  Coprobacter unclassified S1198 GCA 900066575 unclassified S1275  Unclassified unclassified S1330  Enteroscipio rubneri  Sanguibacteroides justesenii  Allisonella unclassified S1349  Raoultibacter unclassified S1185  Asteroleplasma unclassified S1227  Anaerotruncus colihominis  Cloacibacillus unclassified S1375  Candidatus Soleaferrea unclassified S1316  Prevotella colorans |
