## Supplemental Table S4 for "Stability of Oral and Fecal Microbiome at Room Temperature: Impact on Diversity"

**Additional Table S4** Relative abundance and percentage of change in 5 days and 95% confidence interval of top 5 phyla for each method

| OW-CH | | | |
| --- | --- | --- | --- |
| Phylum | **Relative abundance %** | **% Change 5 days** | **95%CI** |
| Actinobacteriota | 16.59 | -7.18 | (-13.14; -1.22) |
| Bacteroidota | 23.92 | 5.58 | (-3.95; 15.13) |
| Campylobacterota | 3.45 | 13.26 | (-1.27; 27.79) |
| Firmicutes | 46.76 | 2.06 | (-2.73; 6.84) |
| Proteobacteria | 7.92 | -6.13 | (-14.28; 2.01) |
| Other | 1.36 | - | - |
| FIT | | | |
| Phylum | **Relative abundance %** | **% Change 5 days** | **95%CI** |
| Actinobacteriota | 24.57 | 4.19 | (-9.56; 17.95) |
| Bacteroidota | 35.29 | -0.96 | (-9.97; 8.05) |
| Desulfobacterota | 2.15 | 24.60 | (-25.82; 75.03) |
| Firmicutes | 28.52 | 0.32 | (-7.08; 7.72) |
| Proteobacteria | 3.51 | -4.02 | ( -28.35; 20.31) |
| Other | 5.96 | - | - |
| ETHANOL | | | |
| Phylum | **Relative abundance %** | **% Change 5 days** | **95%CI** |
| Actinobacteriota | 28.89 | 12.18 | (-3.14; 27.50) |
| Bacteroidota | 27.70 | 1.09 | (-12.19; 14.36) |
| Fusobacteriota | 1.25 | -19.7 | ( -36.36; -3.15) |
| Firmicutes | 35.72 | -2.17 | ( -16.92; 12.57) |
| Proteobacteria | 3.42 | 13.97 | (-33.50; 61.45) |
| Other | 3.02 | - | - |
