## Supplemental Table S5 for "Stability of Oral and Fecal Microbiome at Room Temperature: Impact on Diversity"

**Additional Table S5** Relative abundance, percentage of change in 5 days and 95% confidence intervals of top 5 phyla for each method

| OW-CH | | | |
| --- | --- | --- | --- |
| Order | **Relative abundance %** | **% Change 5 days** | **95% CI** |
| Peptostreptococcales-Tissierellales | 28.49 | -3.52 | ( -11.79; 4.74) |
| Bacteroidales | 19.36 | 4.60 | (-6.71; 15.92) |
| Actinomycetales | 10.86 | -10.75 | ( -18.70; -2.80) |
| Lachnospirales | 6.80 | 8.83 | (2.84; 14.83) |
| Oscillospirales | 6.34 | 25.26 | (3.59; 46.92) |
| Enterobacterales | 4.57 | 2.01 | (-17.80; 21.82) |
| Flavobacteriales | 4.49 | 9.02 | (-2.08; 20.12) |
| Campylobacterales | 3.45 | 13.26 | (-1.27; 27.79) |
| Coriobacteriales | 3.33 | 4.13 | (-7.00; 15.27) |
| Lactobacillales | 2.81 | 0.66 | (-11.55; 12.88) |
| Burkholderiales | 2.51 | -16.73 | ( -26.65; -6.81) |
| Bifidobacteriales | 2.21 | -17.21 | (-25.13; -9.29) |
| Erysipelotrichales | 1.61 | 19.15 | (7.16; 31.15) |
| Pseudomonadales | 0.85 | 0.45 | (-18.96; 19.87) |
| Fusobacteriales | 0.59 | 70.42 | (23.62; 117.22) |
| Veillonellales-Selenomonadales | 0.42 | 24.57 | (-0.44; 49.58) |
| Synergistales | 0.35 | -25.03 | (-62.16; -22.58) |
| Spirochaetales | 0.31 | -42.37 | (-36.47; 18.08) |
| Mycoplasmatales | 0.25 | 21.15 | (-12.68; 54.76) |
| Propionibacteriales | 0.18 | 3.21 | (-9.82; 3.43) |
| Other | 0.23 | - | - |
| FIT | | | |
| Order | **Relative abundance %** | **% Change 5 days** | **95% CI** |
| Peptostreptococcales-Tissierellales | 6.61 | -4.09 | (-20.16; 11.97) |
| Bacteroidales | 33.75 | -0.17 | (-9.98; 9.64) |
| Actinomycetales | 13.28 | 35.55 | (17.57; 53.52) |
| Lachnospirales | 13.37 | 6.61 | (-4.23; 17.46) |
| Oscillospirales | 2.01 | 25.28 | (-22.61; 73.17) |
| Enterobacterales | 2.38 | 3.85 | ( -25.51; 33.21) |
| Flavobacteriales | 1.54 | -8.55 | (-23.39; 6.29) |
| Campylobacterales | 1.45 | 2.88 | ( -19.88; 25.64) |
| Coriobacteriales | 4.17 | 3.28 | ( -15.91; 22.48) |
| Lactobacillales | 1.49 | -8.95 | ( -29.90; 11.99) |
| Burkholderiales | 1.13 | -10.56 | (-33.27; 12.14) |
| Bifidobacteriales | 5.82 | 0.54 | (-29.17; 30.25) |
| Erysipelotrichales | 2.25 | -16.71 | (-35.57; 2.14) |
| Verrucomicrobiales | 2.11 | 25.19 | (-0.53; 50.93) |
| Fusobacteriales | 1.36 | -12.19 | (-31.33; 6.95) |
| Veillonellales-Selenomonadales | 2.37 | 0.882 | (-17.22; 18.98) |
| Synergistales | 0.45 | 6.01 | (-9.40; 21.43) |
| Spirochaetales | 0.43 | -12.99 | (-42.33; 16.34) |
| Corynebacteriales | 0.84 | -18.25 | (-50.09; 13.59) |
| Desulfovibrionales | 2.15 | 24.60 | (-25.82; 75.03) |
| Other | 0.01 | - | - |
| ETHANOL | | | |
| Order | **Relative abundance %** | **% Change 5 days** | **95% CI** |
| Peptostreptococcales-Tissierellales | 9.83 | 9.99 | ( -24.19; 44.19) |
| Bacteroidales | 25.78 | 0.04 | (-14.12; 14.20) |
| Actinomycetales | 18.50 | 9.75 | (-14.67; 34.19) |
| Lachnospirales | 15.95 | 1.04 | (-23.94; 26.03) |
| Oscillospirales | 3.17 | -6.15 | (-33.55; 21.23) |
| Enterobacterales | 1.88 | 7.05 | (-35.16; 49.28) |
| Flavobacteriales | 1.92 | 18.26 | (-8.64; 45.17) |
| Campylobacterales | 1.05 | 2.60 | (-1.82; 7.03) |
| Coriobacteriales | 5.15 | 82.94 | (-103.23; 269.13) |
| Lactobacillales | 1.46 | 35.93 | ( -38.26; 110.13) |
| Burkholderiales | 1.54 | 30.09 | (-26.59; 86.78) |
| Bifidobacteriales | 3.55 | 13.77 | (-13.27; 40.81) |
| Erysipelotrichales | 2.21 | -4.30 | (-24.16; 15.56) |
| Verrucomicrobiales | 0.37 | 140.09 | ( -112.81; 393.0) |
| Fusobacteriales | 1.25 | -19.76 | (-36.36; -3.15) |
| Veillonellales-Selenomonadales | 2.27 | -28.90 | (-54.20; -3.60) |
| Synergistales | 0.48 | 5.39 | (-5.72; 16.51) |
| Spirochaetales | 0.70 | 12.68 | (-16.03; 41.39) |
| Corynebacteriales | 1.06 | 36.68 | (-24.03; 97.40) |
| Staphylococcales | 0.38 | -11.65 | (-30.95; 7.65) |
| Other | 0.01 | - | - |
